# Annotation-Free Prediction of Cancer Cells and Glands and Spatial Analysis of Immune Cells

**DOI:** 10.1101/2025.11.09.687528

**Authors:** Kyeong Joo Jung, Soumya Ghose, Sanghee Cho, Elizabeth McDonough, Chrystal Chadwick, Robert West, James D. Brooks, Dongjun Chung, Fiona Ginty, Raghu Machiraju, Parag Mallick

## Abstract

Prostate cancer is classified as “immune-cold” due to limited infiltration of immune cells and no clear correlation between immune cells and clinical outcomes. However, immune cells are found in prostate cancers and the spatial relationships between these immune cells and cancer cells/glands have not been investigated, partly due to a lack of automated tools that classify both cancerous cells/glands. In this paper, we have developed an end-to-end tool (TOPAZ: **T**issue **O**rganization identification using s**PA**tial proteomics) that combines multiplexed single-cell protein data with histology images to: 1) predict cancerous versus non-cancerous epithelial cells using a Gaussian-mixture model; 2) predict cancerous/non-cancerous gland type using a principal curve estimation. Using TOPAZ to assign cancer and non-cancerous labels to cells and glands, we extracted multiscale spatial features from the classification results—including immune dense-region geometrical features and cell-to-gland distances— and correlated the features with risk of biochemical recurrence and cancer grade. Tissue-microarrays containing 753 cores from 217 prostate cancer patients underwent multiplexed immunofluorescent imaging (Cell DIVE, Leica) for epithelial cell markers (panCK26, S6, NaKATPase), basal cell markers (p63, CK5), a cancer cell marker (AMACR), and T cell markers (CD3, CD4, CD8, FOXP3, CD68). Cancerous/non-cancerous cell classification from TOPAZ achieved 82% sensitivity and 99% specificity against expert annotation, and the pipeline further predicted cancerous/non-cancerous glands without manual threshold tuning. Regulatory-T-cell and helper-T-cell percentages decreased, and macrophage percentage increased with grade increase (P < 0.05). When the median distance from cancerous gland centroids to the nearest regulatory or helper T-cell exceeded approximately 50 µm, the hazard of biochemical recurrence doubled (log-rank P < 0.01). The open-source Shiny app TOPAZ (https://chunglab.bmi.osumc.edu/TOPAZ) packages the workflow, predicting individual cell types and gland shapes. By combining probabilistic cell typing with gland-shape modeling, TOPAZ yields interpretable multiscale spatial features linked to prognosis and is released as an open web app for unrestricted use.

**Author Summary:** Spatial distribution of cancerous cells/glands and immune cell distribution has not been considered as a prognosticator in prostate cancer. In addition, automated tools that can quantify and integrate these distributions are lacking. We combined high-dimensional single-cell protein measurements with histology images to map gland structure in prostate cancer tissue. Our web-based tool, TOPAZ (https://chunglab.bmi.osumc.edu/TOPAZ) predicts whether each epithelial cell and gland in virtual H&E image is cancerous or not. Once the predictions are made, a spatial analysis workflow helps quantify spatial features of immune cells relative to the glands and correlate with recurrence risk and grades. Across 753 tissue cores from 217 prostate cancer patients, helper and regulatory-T-cells located more than about 50 µm away from cancerous epithelial glands were associated with a higher risk of biochemical recurrence. The pipeline provides new insights for researchers and pathologists into prostate cancer progression and biochemical recurrence through integration of spatial location of cancer glands and immune cells.

## Introduction

Prostate cancer is the second leading cause of cancer and fifth leading cause of death in men globally [1,2]. In the US, it is the most common cancer and the second leading cause of death (255,395 new cases and 33,881 deaths in 2022 [3]). Gleason Score [4] is the most commonly used histology-based method for assigning diagnostic grades based on glandular morphologic patterns in excised tissue and range from 1 to 5. Since a patient tissue can be assigned to multiple patterns, Dr. Gleason proposed that the scores of the most common and second most common patterns be summed [5], and are now summarized as a Grade Group. The Gleason Score (sum ranging from 2-10) is an important predictor of patient prognosis (recurrence, overall survival, etc.) whereby a Gleason Score less than or equal to 3+3=6 (Grade Group 1) typically indicates cancers that exhibit indolent behavior, while Grade Groups 2 (3+4=7), 3 (4+3=7), 4 (4+4=8) and 5 (4+5=9 and above) exhibit progressively increasing risks of poor outcomes [6]. Simultaneously, there are a growing number of prognostic tests for prostate cancer including the Decipher mRNA test [7]. This test provides risk prediction and is used clinically in the selection of hormonal therapy in men undergoing radiation for localized prostate cancer. However, these tests are based on bulk mRNA analysis [8] and do not provide information on how spatial variations in the tissue structure correlate with underlying mechanisms of the initiation, growth, and maintenance of prostate cancer.

It is crucial to understand disease mechanisms during cancer progression and the factors that contribute to clinical outcomes. This involves examining the changes in cell types (cancerous epithelial, non-cancerous epithelial, immune, and stromal), their spatial distribution, and their potential interactions based on proximity. Recent advances in cancer systems biology have demonstrated that transcriptomic, proteomic, and structural heterogeneity as manifest by glands and the tumor micro-environment (cellular behavior/functions near tumor, immune cells, blood vessels, etc.) are key drivers of disease progression [9–12]. However, bulk RNA-sequencing disregards critical information about cellular interactions within the tumor microenvironment. Recent advancements in single cell imaging platforms now allow spatial analysis of *in situ* cellular expression of dozens of proteins and thousands of transcripts at single cell level, which has accelerated detailed analysis of cellular interactions and distributions in intact tissues [13].

However, existing pipelines do not fully incorporate spatial information of morphological structures or analyze multi-scale factors impacting tissues-at-large beyond individual cells and often require consideration of glandular structures and spatial arrangements within tissues. It also includes cellular interactions, such as immune cell infiltrations and their interactions with cancerous cells. In prostate cancer, disease progression is strongly correlated with variations in the spatial architecture of glands, which is formally captured by the Gleason scoring system [4]. Despite the well-studied impact of the changes in glandular morphology, the interactions between various tissue structures such as immune cells, fibroblast, blood vessels, and hypoxic regions remain unexplored.

Existing computational methods fall short in modeling complex tissue architectures. For example, MAPS [14], ASTIR [15], CellSighter [16] use probabilistic modelling, deep recognition neural networks, and convolutional neural network (CNN) respectively to classify distinct cell populations based on protein expression profiles. While these methods capture spatial context based on cellular locations, they do not integrate spatial cellular data with tissue morphology, nor do they consider spatial organization of other cellular actors including salient immune cells.

To address these challenges, we have developed a computational pipeline that classifies cancerous epithelial (CE)/non-cancerous epithelial (N-CE) cell and gland types and analyzed their spatial organization across all grades in prostate cancer. Our pipeline unifies single-cell proteomics with virtual H&E (vH&E) to resolve CE architecture and its immune context and analyze the results in association with the clinical outcomes. This pipeline provides three major novelties: i) **TOPAZ** (**T**issue **O**rganization identification using s**PA**tial proteomics, an open access application [17]) that performs **annotation-free classification** of CE and N-CE cells/glands using Gaussian mixture model (GMM) and principal curve estimation. This removes the need for manual thresholding and enables classification through large patient cohorts; ii) We introduce **shape predictions** which enable us to predict gland boundaries and quantify spatial features such as area and distances which prior methods did not focus on; iii) After the classification, we investigate **CE gland-immune interactions** through infiltration events and associate it with the clinical outcomes such as the grade groups, and biochemical recurrence.

Beyond these novelties, pipeline implementation can potentially **generate labels for the deep learning models** which can lead to automated tissue classification without manual expert annotation. Finally, we demonstrated the utility of the framework by comparing **AMACR protein expressions in classified CE and N-CE cells** across prostate cancer grades and **spatial distribution analysis** to identify glandular geometrical features and associated with clinical outcomes. In summary, these innovations allow us to move from cellular-level classification to glandular-level morphology and gland-immune-cell interactions revealing desired multi-scale factors of prostate cancer progression.

**Fig 1** illustrates our pipeline for classifying CE and N-CE cells/glands (TOPAZ pipeline), performing spatial distribution analysis, and predicting biochemical recurrence based on extracted spatial features. The pipeline ingests virtual H&E images, single-cell data for each protein biomarker, and gland masks (see Materials and Methods).

**Fig 1.**
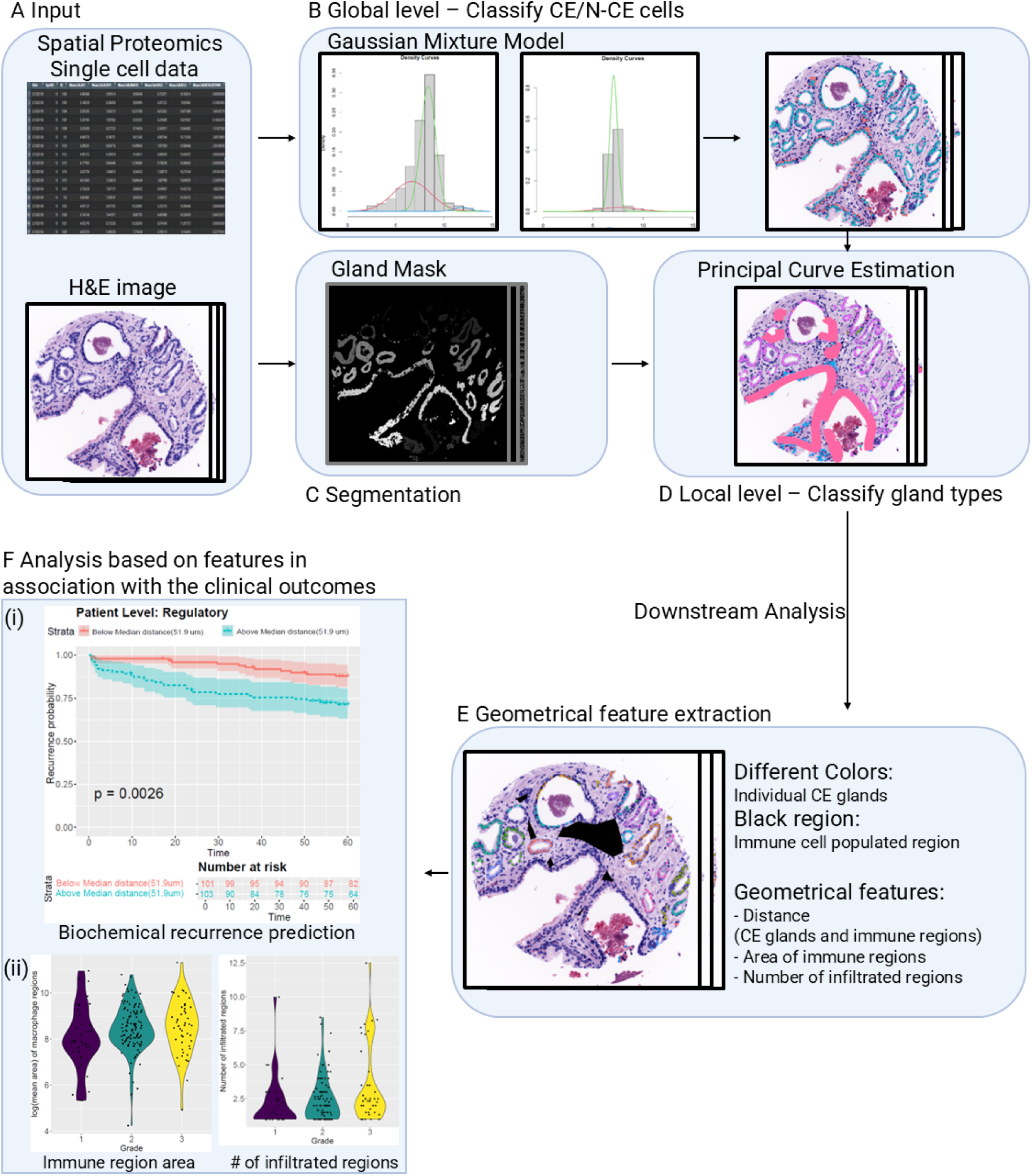
Overview of the pipeline integrating single cell protein, histology images, and clinical outcome analysis. **(A) Input Data:** The pipeline begins with multiplexed immunofluorescence single-cell protein data and virtual H&E images as inputs; **(B) Global Level Analysis:** A Gaussian Mixture Model (GMM) was applied to classify N-CE, CE cells based on the distribution of single-cell expression of p63 and CK5. The resulting probability maps were overlaid onto virtual H&E images for visualization. **(C) Segmentation - Gland Mask:** A gland segmentation mask was generated to identify individual glandular structures within the tissue via combination of pretrained DL model and GMM; **(D) Local Level Analysis - Principal Curve Estimation of glands:** Within each segmented gland per tissue, principal curve estimation was applied to capture the intrinsic gland shape and assign cancer/non-cancerous epithelial (CE/N-CE) cell labels based on their spatial proximity to the estimated latent curve; **(E) Geometrical feature extraction:** Based on the predicted results, geometrical features such as distances between CE glands and immune regions, area of immune regions, and number of infiltrated regions are extracted; **(F) Association with Clinical Outcomes:** The final output involved extracted feature-based analyses associating spatially classified glands with clinical outcomes. This included **(i) Biochemical recurrence prediction** to correlate distance between CE and immune regions at patient level; and **(ii) Violin plots** to visualize the features such as area, number of infiltrated regions across grades.

In the following sections, we first describe the patient cohort, tissue microarray construction, and multiplexed imaging platform used in this study (Materials and Methods). Next, we introduce the TOPAZ pipeline which performs annotation-free classification of CE/N-CE cells and glands. We then extract spatial features describing CE-immune interactions. Next, we present results on CE/N-CE/stroma and immune cell compositions, AMACR marker comparison, spatial associations with the clinical outcomes, and software implementation for classification. We conclude by discussing the implications of these findings for prostate cancer biology and by outlining potential applications of the pipeline.

## Materials and Methods

### Patient cohorts

Of tissue microarrays (TMA) comprising 754 cores from 234 prostate patients, QC exclusions yielded 217 patients and 753 cores for analysis including 42 patients who had 5-year biochemical-recurrence data. Analysis was conducted on prostate cancer patients from the Stanford Urology Tissue Bank. Samples were collected under IRB-approved protocols (IRB #11612) from more than 600 men who were diagnosed with prostate cancer between 1996 and 2009 and treated by radical prostatectomy at Stanford University.

### TMA construction

Gleason scoring was made based on examination of H&E-stained tissue sections of the entire radical prostatectomy specimen by board certified pathologists with special expertise in genitourinary pathology. Four cores of cancer tissue were sampled per specimens and linked to a secure database including pathology reports, tumor measurements, patient age, pre-operative prostate-specific antigen (PSA) levels, clinical stage, and follow-up information for biochemical recurrence. Summary statistics for demographics are shown in Table A in S1 Text. Although data from 217 patients were used for the classification task, two patients without biochemical-recurrence information were excluded from the analysis involving patient outcome. These samples formed the foundation for subsequent multiplexed imaging and spatial proteomic analyses.

### Multiplexed imaging

TMAs underwent imaging using the Cell DIVE platform (Leica Microsystems) [18] which provided multiplexed imaging of proteins that had been previously reported to be associated with prostate cancer diagnosis, progression and immune response. For the purpose of this study, we focused on a subset of 12 markers including basal cell markers (CK5, p63); epithelial/stroma segmentation markers (NaKATPase, S6, panCK-PCK26, panCK-AE1); nuclear stain (DAPI); immune cell markers (CD3, CD4, CD8, CD68, FOXP3); and cancerous cell marker (AMACR).

Prior to the multiplexed imaging, all antibodies were subjected to a standardized characterization protocol [19]. Multiple clones for each target were screened for sensitivity and specificity using serial sections of a commercially available multi-tissue microarray (MTU391, Pantomics) or a custom prostate tissue microarray and then evaluated for epitope stability to dye inactivation. The best performing clone was then conjugated to a fluorescent dye and re-evaluated. Sequential staining and dye inactivation of the study TMAs was performed as described previously [18]. In brief, samples underwent deparaffinization, antigen retrieval, and blocking, followed by iterative staining and imaging on a commercial Cell DIVE imager (Leica Microsystems) at GE HealthCare Technology & Innovation Center (HTIC). DAPI and autofluorescence were imaged in every round and images underwent automated post-processing including illumination and distortion correction, image registration, and autofluorescence signal subtraction. The combination of DAPI and autofluorescence images was also converted to a pseudo-colored H&E stain (virtual H&E), which was used for pathology review and gland analysis. Processed images were then segmented to generate single-cell datasets for quantitative analysis.

### Single-cell data generation

A customized Fiji (ImageJ) plug-in was used for single-cell data generation [20]. Cells in the epithelial and stromal compartments were segmented using DAPI and pan-cytokeratin, while S6 and NaKATPase were used for subcellular analysis of epithelial cells. Each segmented cell was assigned an individual ID and spatial coordinate, enabling subsequent cell-to-cell interaction analyses. Markers were quantified in each compartment and the entire cell. Our TIFF image format allowed us to use open-source tools such as QuPath for image review and annotation [21]. The cell table (containing IDs, coordinates, and intensities) served as input to QC/normalization and then to the classification/analytical pipeline summarized below.

### Cell-level QC and normalization

Images from each tissue core were reviewed for tissue quality to be included in the analysis—including tissue loss, damage, and segmentation quality. Several QC were then applied at the cell level:

1. Nuclear integrity—epithelial cells were required to include one clearly segmented nucleus but no more than two within each sub-compartment.
2. Sub-compartment areas (epithelial): each sub-compartment (nucleus, membrane, cytoplasm) for epithelial cells was required to have area greater than 10 pixels, but not more than 1500 pixels.
3. Whole-cell area: the total cell area was required to be between 50–3,500 pixels for epithelial cells and 30–1,500 for stromal cells.
4. Cyclic imaging stability: Using the correlation of cell level DAPI signal from each staining and imaging round, a quality score was generated for every cell in each image, which ranges from 0–1: 0 being no registration and up to 1 for perfect registration. Only cells with quality scores above 0.85 were included in the statistical analysis. Scores below 0.5 are generally due to tissue shifting/movement and loss.
5. Signal review: intensity outliers were visually inspected per biomarker and excluded if artifactual.
6. Normalization [22] minimized batch effects, and log₂ transformation addressed skewed intensity distributions.
7. Marker-specific QC: epithelial cells not passing QC (Cyclic imaging stability) for p63 or CK5 were filtered out before CE/N-CE classification.

After QC, we analyzed 724,742 epithelial cells and 552,702 stromal cells. Sample vH&E and multiplexed p63/CK5 images are shown in Fig A in S1 Text. Summary statistics of the single-cell data appear in Table B in S1 Text.

### Immune cell labeling

After establishing epithelial and stromal QC criteria, immune phenotyping was performed using the Cell Auto Training (CAT) model [23]. Previously published CAT model was applied to immune cell markers (CD3, CD4, CD8, CD68, and FOXP3) to generate binary positive/negative labels for each marker at single-cell level. By the cell type definition of T cells and macrophages, T-helper cells (T_H_: CD3⁺CD4⁺), T-regulatory cells (T_reg_: CD3⁺CD4⁺FOXP3⁺), and macrophages (CD68⁺) were labeled. These designations were used to quantify immune-cell proximities to cancerous glands in downstream analyses.

### Gland mask segmentation

We developed a hybrid unsupervised, self-supervised framework for gland-mask segmentation. A pretrained deep-learning (DL) nuclei model [24] handled DAPI segmentation; unsupervised multi-class GMM classified epithelium via CK8/18 images. Fusing both masks yielded probabilistic gland masks and was further refined using the pretrained DINOv2 model [25] in a frozen feature-extraction mode to remove spurious regions. As observed in Fig 2, gland segmentation was correctly performed by our model, opening opportunities for further classification and downstream analysis.

**Fig 2.**
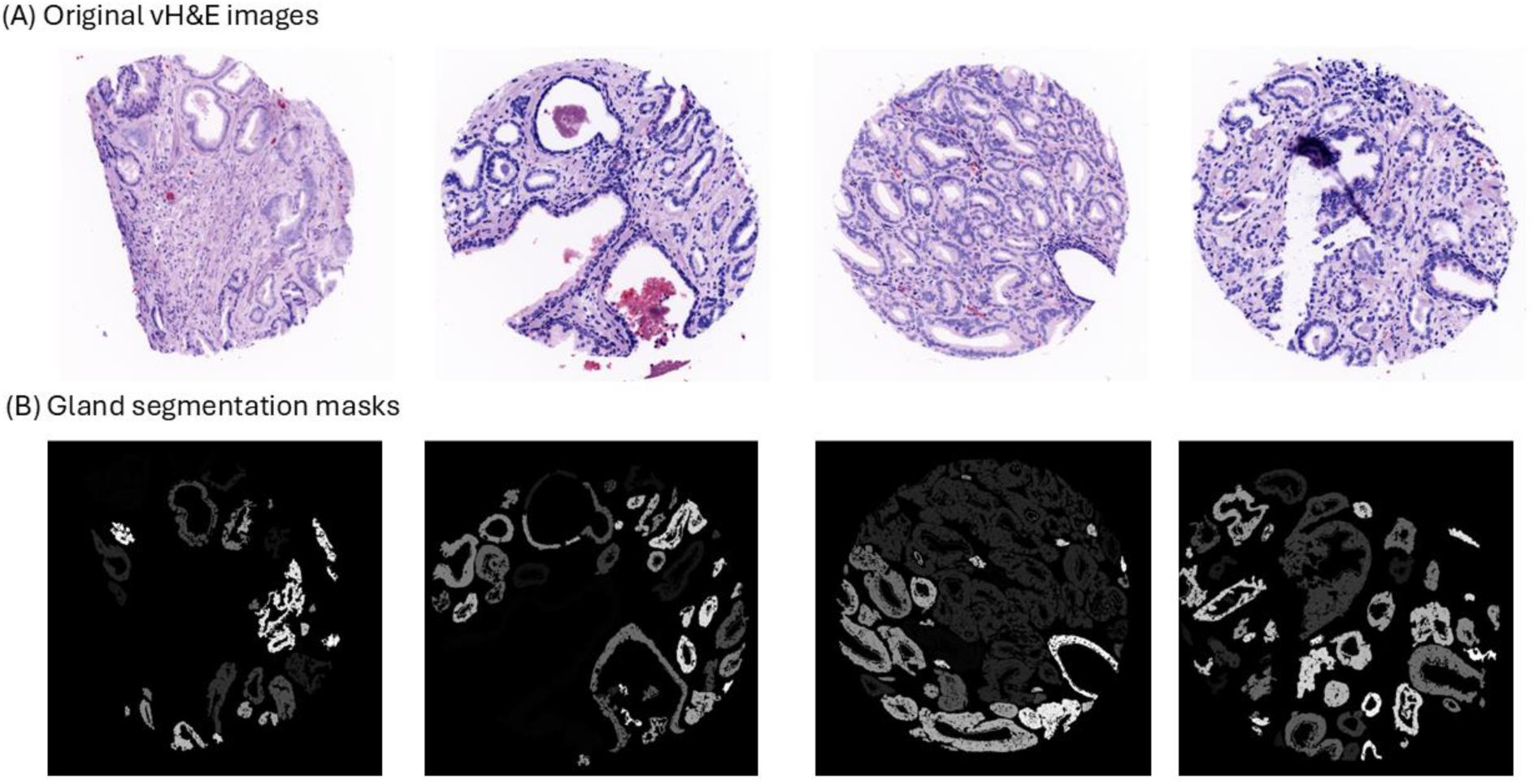
Examples of gland segmentation masks. (A) 4 different prostate virtual H&E (vH&E) images. (B) 4 different gland segmentation masks of vH&E images. Same intensity (color) indicates same individual glands.

### Overview of classification/analytical pipeline

The classification process (TOPAZ pipeline) comprised three steps (Fig 3):

1. Identification of CE versus N-CE cells via Gaussian Mixture Model (GMM), where CE cells were negative for both markers (p63⁻, CK5⁻) and N-CE cells positive for ≥ 1 marker (p63⁺ or CK5⁺).
2. Integration of gland masks and prediction of cell types using principal-curve estimation and a minimum-spanning tree (MST) to preserve gland continuity.
3. Capture of gland shape and assignment of CE/N-CE cells by curve expansion.

**Fig 3.**
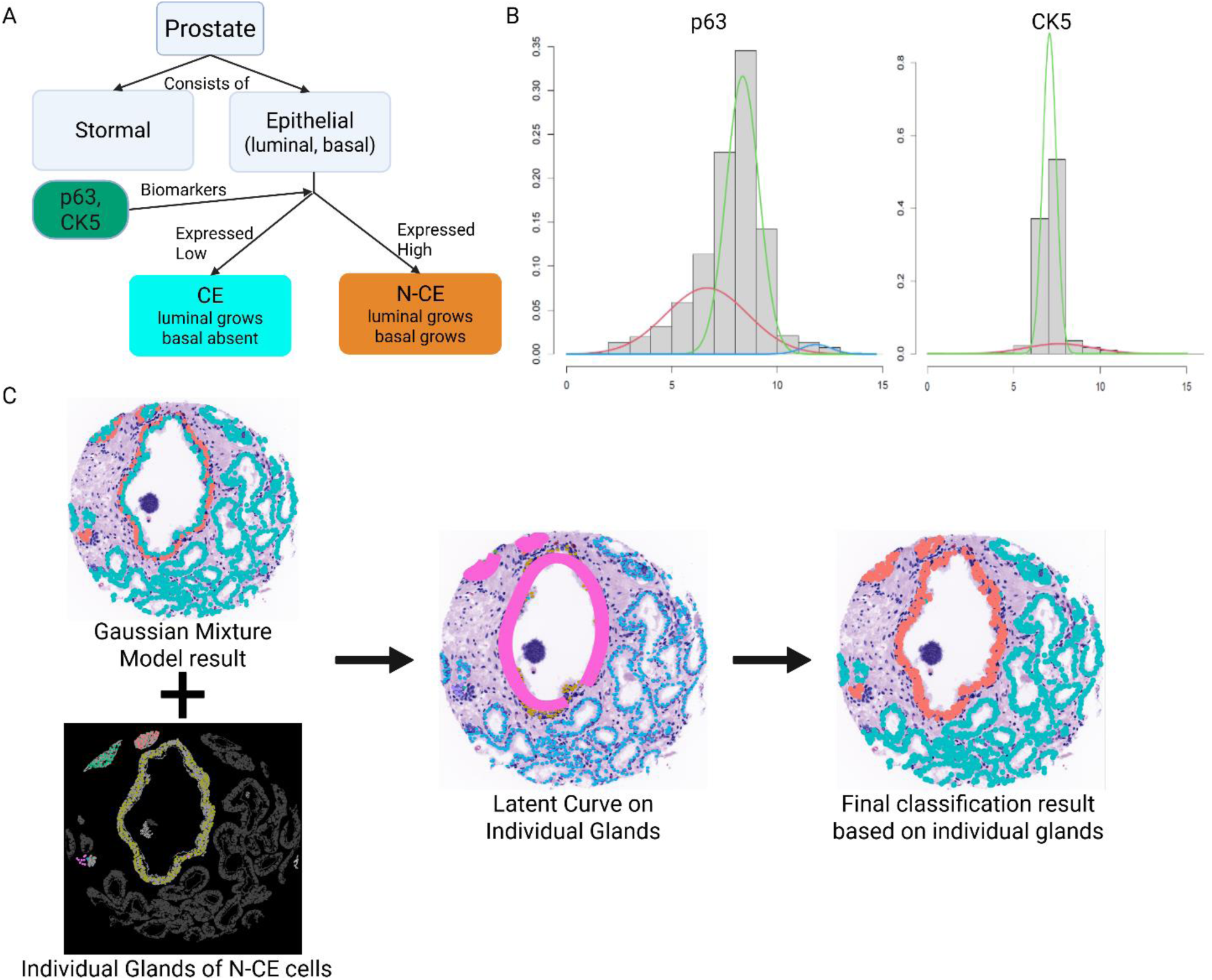
(A) Decision tree of prior knowledge biomarkers of a cell type (CE / N-CE), (B) Gaussian mixture components of two different biomarkers (p63, CK5), (C) By combining GMM result (upper left) and gland masks, we generated latent curve on the individual glands for N-CE cells. Lastly, we assigned the groups of the cells within certain distances from the curve (band) as the same cell type.

This order reflected the hierarchical organization of tissue structure, from cells to glands. Feature extraction then defined geometric and infiltration metrics, and statistical analyses evaluated their association with grade and recurrence. This multi-step pipeline integrates spatial proteomics and morphometric analysis for patient-level inference

### Cancerous epithelial (CE) and non-cancerous epithelial (N-CE) cell identification

We applied a Gaussian Mixture Model (GMM) [26] using Expectation-Maximization (EM) [27] to classify CE and N-CE cells. The EM algorithm estimated mixture parameters distinguishing foreground (positive expression) and background (negative expression) distributions for p63 and CK5 (Fig 3B). Unlike prior pixel-level approaches [24], our implementation operated at the single-cell level for improved biological interpretability. For each biomarker, the mixture model is defined as:

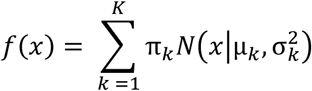

where *N* denotes the normal distribution, π_*k*_ are the mixing proportions such that 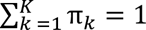, and 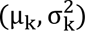 are the means and variances of the *k*-th component distribution for each biomarker. Here, *K* represents the number of mixture components. In our workflows, we normally use *K* as 2 or 3 based on the histogram because in spatial proteomics, the positive cells are located mostly at the high-end of the histogram.

We set ε = 1e-08 and a maximum of 1,000 iterations. The EM algorithm iteratively computed posterior probabilities 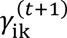 and updated parameters until convergence for each cell *i* and each component *k* at iteration *t+1* for each biomarker in the E-step:

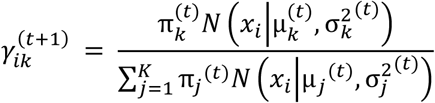

and the M-step, which updates the parameters 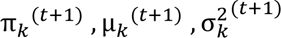

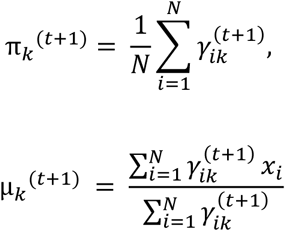

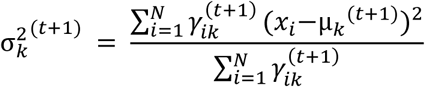

Cells are classified as positive where the expression value has higher posterior probability than other components from the component, which is on the highest side, indicating they belong to the foreground distribution.

Following EM classification, we combined p63 and CK5 positivity using a logical OR. Thus, a cell was considered positive if it is positive for at least one biomarker (p63 and CK5). If *C_i,b_* represents the classification (1 for positive, 0 for negative) of cell *i* for biomarker *b*, the overall positivity *P_i_* is given by:

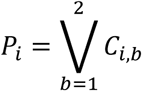

where **Ⅴ** denotes the logical OR operation.

### Cell annotation workflow for validation

Samples representing low and high average p63 intensity were randomly selected across three TMAs (n = 12 cores, 23,208 cells). Two expert biologists (A, B) manually annotated cells using QuPath tools blinded, and coordinates were exported for validation. The inter-rater agreement between the two reviewers was almost perfect (Cohen’s κ = 0.925, p < 0.001), indicating high consistency in manual labeling. Based on manual annotations, model predictions achieved sensitivity of 81.05% (A) and 81.88% (B) with ≈ 99% specificity (Table 1). High specificity suggests robustness of cell-type discrimination across annotators.

**Table 1.** Sensitivity and Specificity with 2 expert annotations.

| Expert | Sensitivity | Specificity |
| --- | --- | --- |
| A | 81.05% | 99.73% |
| B | 81.89% | 99.66% |

### Cancerous/Non-cancerous gland detection using principal-curve estimation

To characterize gland structure, we first imposed a spatial ordering of N-CE cells using a minimum spanning tree (MST) [28] to ensure continuity of gland shape. The MST connected cells based on their pairwise Euclidean distances, producing an ordered path along the gland contour. Using this ordering, we estimated latent curves representing intrinsic gland shapes via principal-curve estimation [29].

Formally, given a set of data points *x_i_*, principal-curve γ(*t*) is defined as the function that minimizes the average squared distance between each data point and its projection on to the curve:

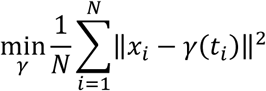

where *x_i_* represents the spatial coordinates of N-CE cells, and *γ*(*t_i_*) are the projections of these points onto the curve. Cubic smoothing splines minimized average squared distances between N-CE cell coordinates and their curve projections. The smoothing level (df = 8) provided a balanced trade-off between fidelity to gland morphology and over-smoothing, consistent with the bias–variance principle of smoothing spline regularization [30].

Latent-curve estimation was applied only to glands meeting a size threshold of 25,000 pixels per N-CE cell count ratio, excluding sparsely populated structures. Thus, this threshold prevents the misclassification of abnormally large glands or sparsely populated N-CE cells as N-CE glands. If a gland is too large compared to the number of N-CE cells detected within it, this suggests that the region may not represent a true N-CE gland, but rather a CE cell-dominated gland with a few scattered N-CE cells. Gland area of 25,000 pixels was empirically chosen based on observed distributions of gland sizes and N-CE cell densities in prostate tissue. By applying this threshold, we ensured that only glands with a reasonable proportion of N-CE cells were used for latent curve estimation, improving the accuracy and biological relevance of the segmentation.

### Expanding the latent curve to capture gland shape

A principal curve alone delineates trajectory but not gland membership. To define gland boundaries, we expanded the curve into a 10-pixel perpendicular band encompassing adjacent cells, assigning those within as N-CE cells. The 10-pixel width was determined empirically based on the typical glandular width observed from the number and size of epithelial cells. Each cell received a label based on proximity, ensuring spatially coherent gland assignments. The same logic was later applied to CE glands. This method improved geometric feature derivation such as gland area and centroid mapping.

### Spatial feature extraction for downstream analysis

Following classification, we computed three geometry-based metrics using R (version 4.3.2.) (Fig 4):

1. *Cell-centric distance* — shortest Euclidean distance (1 µm = 0.325 pixels (the camera pixel size = 6.5 µm, and imaging was performed using a 20× objective).) from each T cell (T_reg_ = CD3⁺CD4⁺FOXP3⁺; T_H_ = CD3⁺CD4⁺) to the centroid of the nearest CE gland.
2. *Region-centric distance* — shortest boundary-to-boundary distance between macrophage-dense (MD) regions (CD68⁺ alpha-shape clusters [31]; description in Fig B in S1 text) and CE gland boundaries.
3. *Infiltration indicator* — distance = 0 when MD regions overlapped glands.

**Fig 4.**
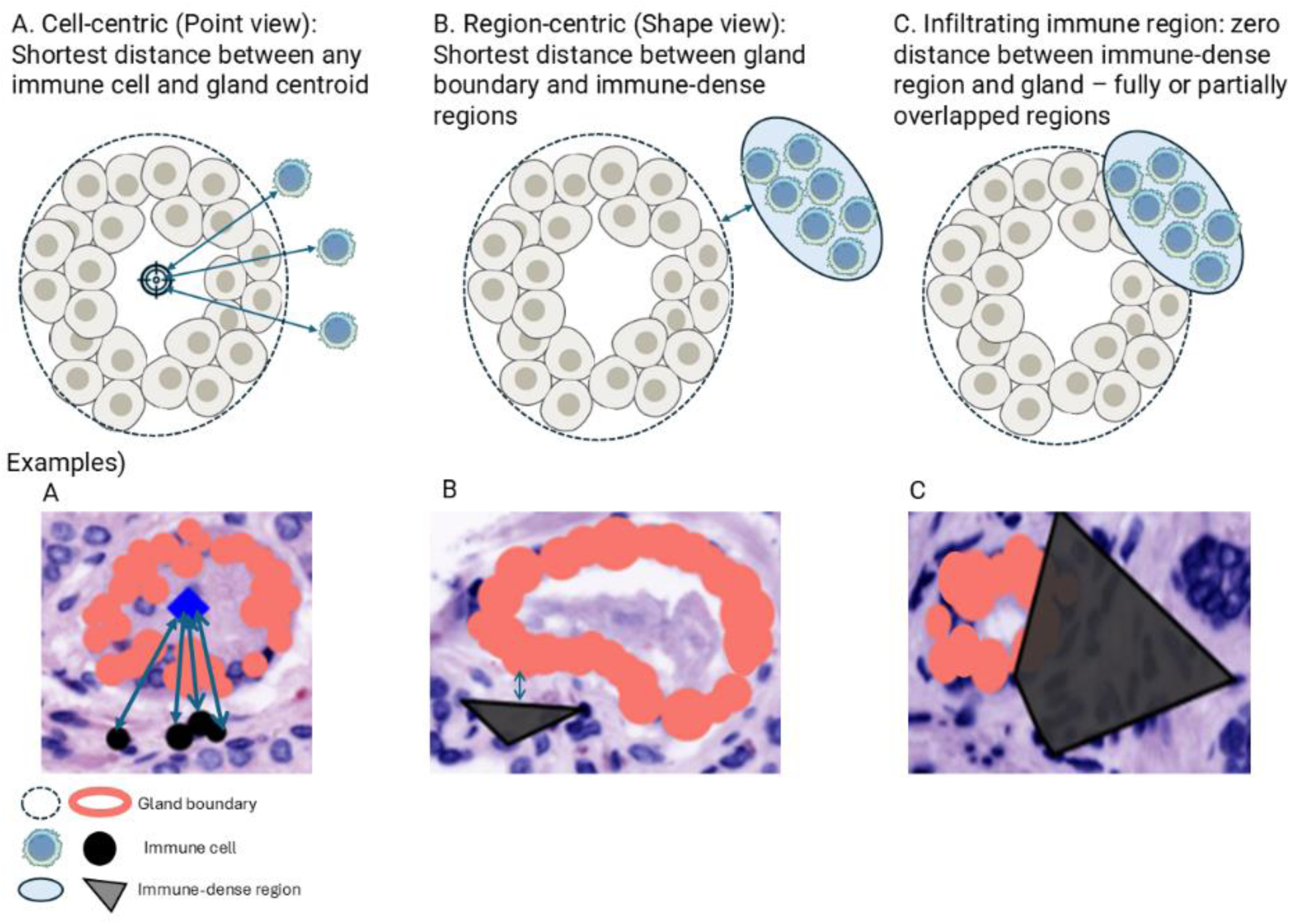
Explanation of the extracted geometrical features from the classified results. (A) Cell-centric (Point view) distance: shortest distance between immune cells and gland centroid. (B) Region-centric (Shape view) distance: shortest distance between gland boundaries and immune-dense regions. (C) Definition of infiltrating immune region: zero distance between immune-dense region and gland – fully or partially overlapped regions.

At the patient level, T-cell distances were summarized by median; MD-region areas by log mean; and infiltration counts tallied per patient. We excluded 37 of 753 tissue spots lacking distinct gland segmentation before statistical analysis.

### Statistical analysis of downstream analysis

Cell-type and compartment compositions across grades were tested using permutation χ² tests with 10,000 label shuffles and Benjamini–Hochberg adjustment [32]. AMACR expression was compared between CE and N-CE cells using two-sided permutation tests within each grade and pooled across grades. Grade trends employed Kruskal–Wallis [33] test. Kaplan–Meier survival analysis with log-rank tests assessed biochemical-recurrence risk, dichotomizing distances at patient-level medians. Tests were two-sided with α = 0.05. All analyses were conducted in R (version 4.3.2) using base and survival packages.

## Results

### Cellular composition and immune prevalence stratified by grade

Using the pipeline and features defined in Methods, we first summarize compositions, evaluate biomarker expression by grade, and then test spatial features for associations with grade and recurrence. Using the TOPAZ pipeline, we determined CE/N-CE cell counts and distributions across different grades. Representative examples of p63/CK5-positive cells and corresponding classified CE/N-CE glands are shown in Fig 5, where orange and blue denote N-CE and CE cells, respectively.

**Fig 5.**
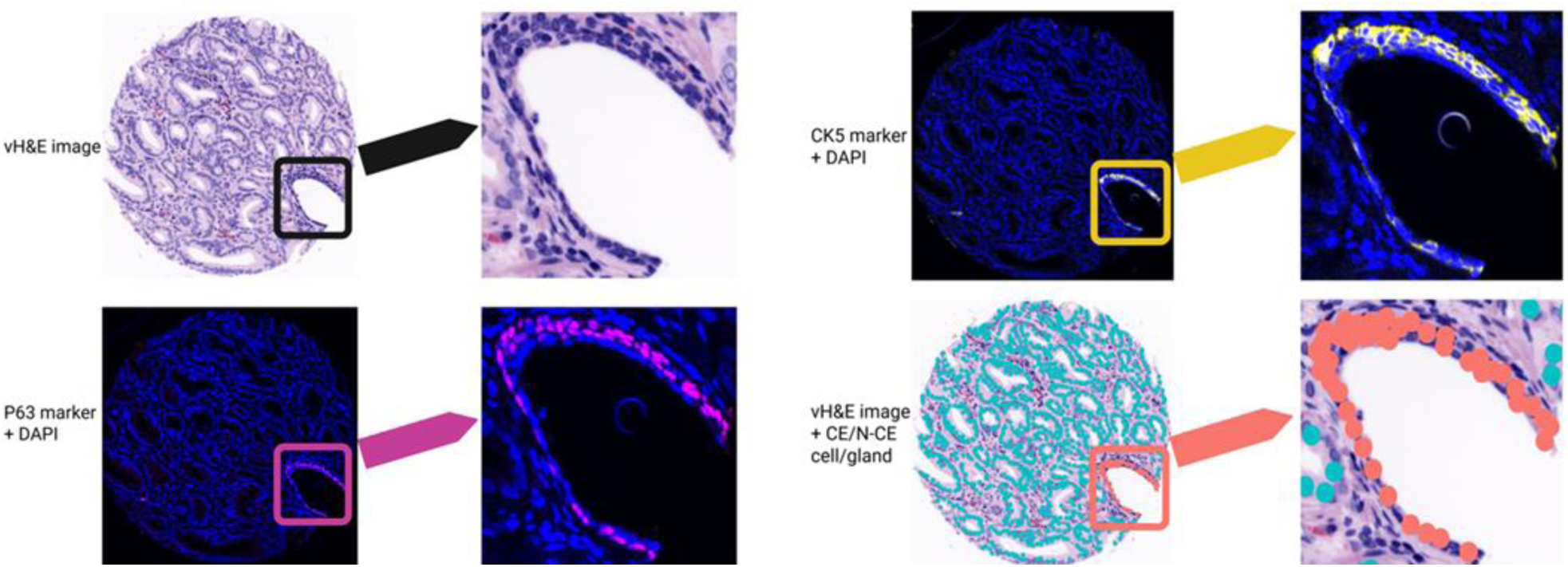
Example of CE (blue) and N-CE (orange) cell/gland identification. Left panel: full-size tissue image. Right panel: zoomed region of interest (ROI) from the same image. Top left: vH&E image (black). Bottom left: p63 + DAPI (magenta). Top right: CK5 + DAPI (yellow). Bottom right: classified cells overlaid on H&E image (orange).

Next, we examined proportional differences among CE, N-CE, and stromal cells, as well as among immune-cell types, across grades. As shown in Fig 6A, CE proportions increased and N-CE decreased with higher grade, while stromal fractions remained relatively constant. A permutation test (see Methods) indicated a significant association between grade and cell-type proportions (p < 0.0001; Fig C in S1 Text). We then evaluated immune-cell prevalence by grade. Fig 6B(i) shows that helper T (T_H_)-cell and macrophage proportions increased with grade (p < 0.001 for each), whereas regulatory T (T_reg_)-cell proportions remained low and stable (p = 0.065). Fig 6B(ii) further partitions immune-cell proportions by compartment (N-CE, CE, stroma). The grade-related increase in T_H_ cells was confined to stroma (p < 0.001).

**Fig 6.**
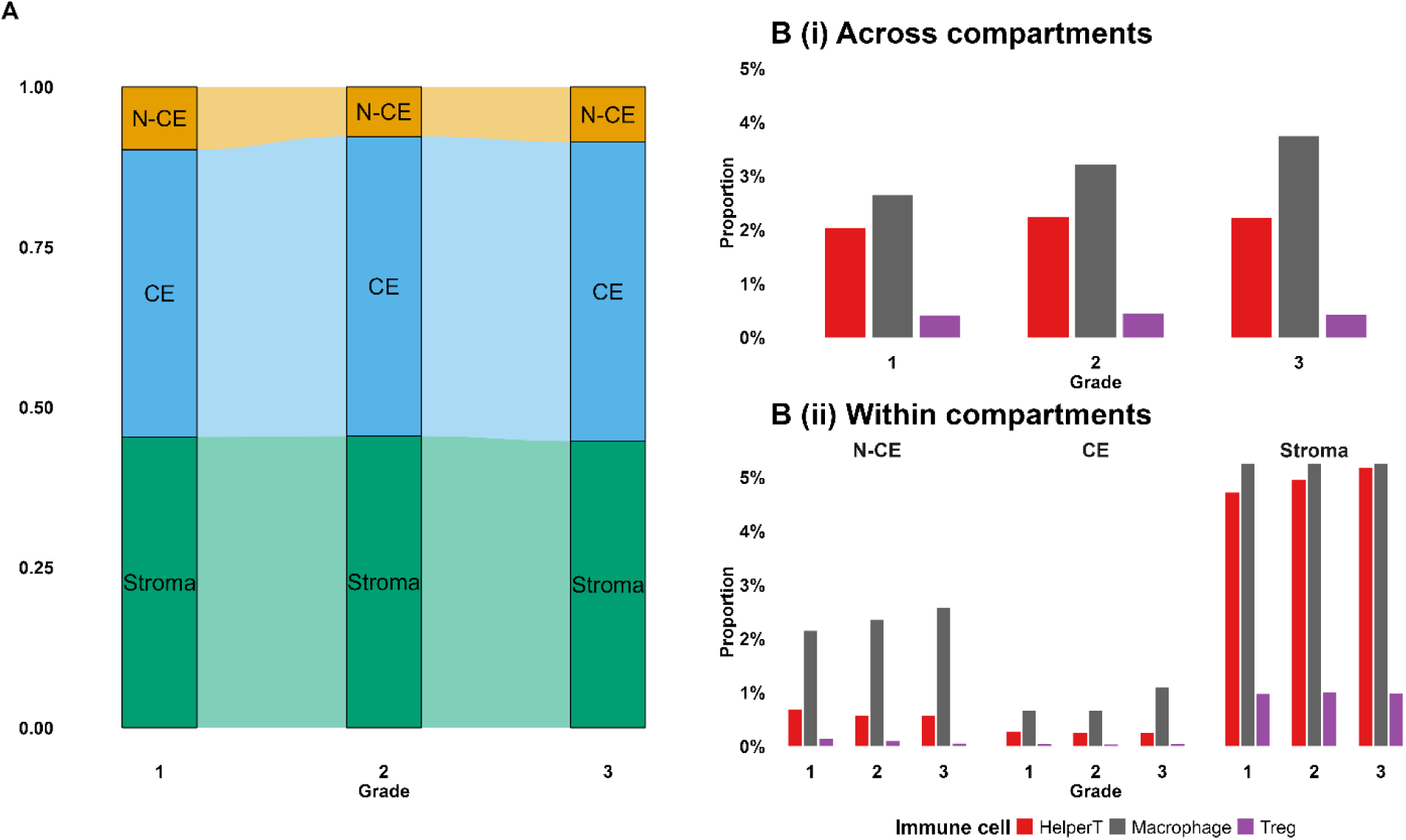
(A) N-CE, CE, and stromal cell distributions by grade. (B) Immune-cell distributions (T_H_, macrophage, T_reg_) across grades: (i) entire sample (N-CE + CE + stroma); (ii) by compartment.

Likewise, macrophage proportions rose with grade within both stromal and epithelial compartments (p < 0.001 and p < 0.05, respectively). These patterns remained significant after multiplicity adjustment (Benjamini–Hochberg; Fig C in S1 Text).

### AMACR expression distinguishes CE from N-CE across grades

AMACR (alpha-methylacyl-CoA racemase), a key enzyme involved in fatty-acid metabolism, serves as a well-established diagnostic biomarker that is markedly overexpressed in prostate-cancer epithelium compared with benign glands [34]. To validate the accuracy of our cell-type classification, we compared AMACR expressions between CE and N-CE cells identified by the TOPAZ workflow. Figure 7 presents median AMACR intensities and log-fold changes across (A) all grades, (B) Grade 1, (C) Grade 2, and (D) Grade 3. A log-fold change greater than 1 indicates higher expression in CE, whereas values below 1 correspond to N-CE expression. Across all grades, CE cells consistently demonstrated elevated AMACR levels, and two-sided permutation tests confirmed these differences as statistically significant (Fig D in S1 Text).

**Fig 7.**
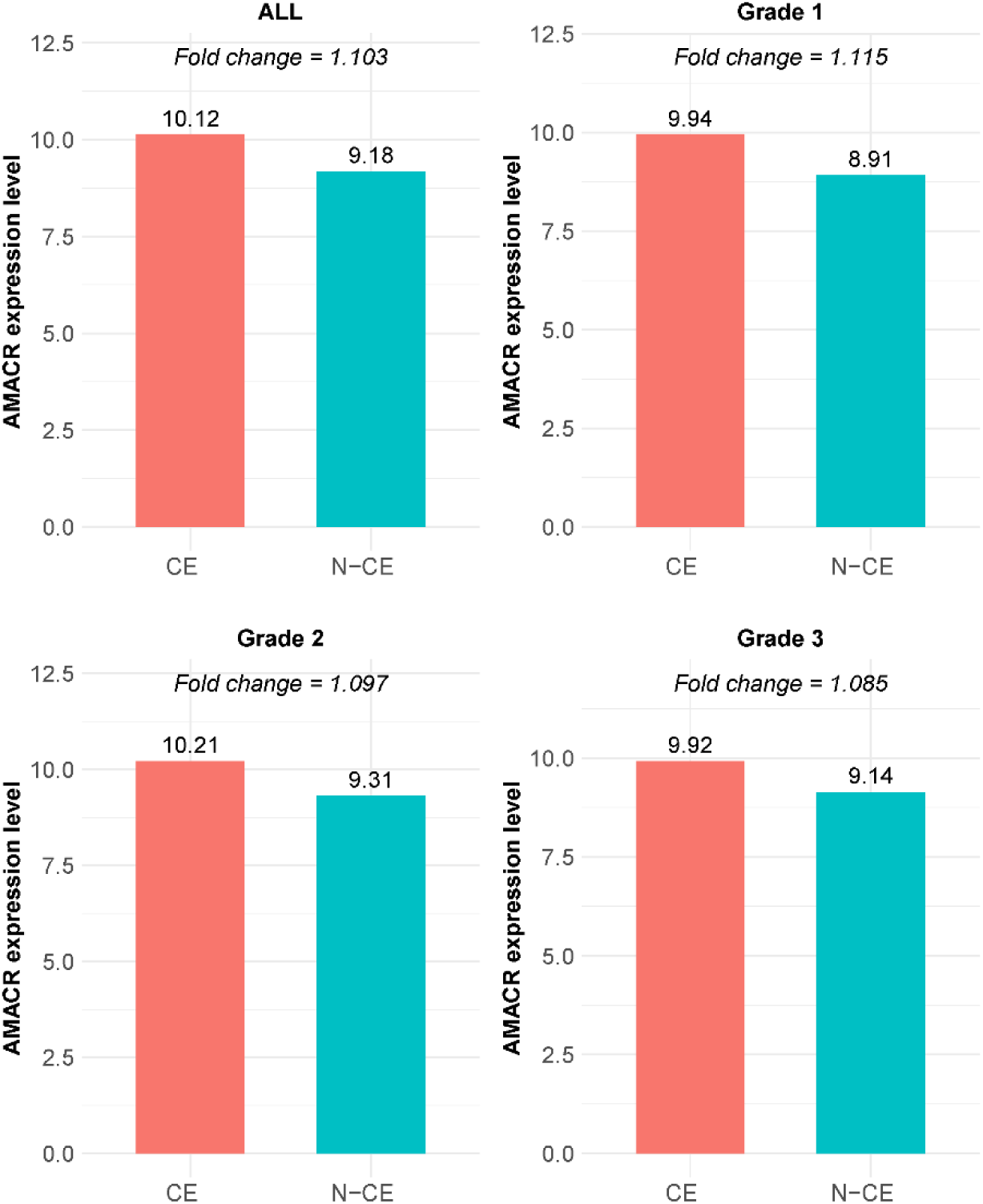
AMACR expression and log-fold changes between CE and N-CE: (A) all grades; (B) Grade 1; (C) Grade 2; (D) Grade 3.

### Spatial distribution of immune cells related to CE glands

We next evaluated geometric features defined in Methods—cell-centric (*point-view*) T-cell distances to the nearest CE-gland centroid, region-centric (*shape-view*) macrophage-dense (MD) region distances to CE boundaries, and infiltration events (distance = 0).

Using the point view, patients with greater T-cell–to-CE distances had increased risk of biochemical recurrence across grades. Dichotomizing by cohort medians (T_reg_ ≥ 51.9 µm; T_H_ ≥ 51.7 µm), Kaplan– Meier curves demonstrated shorter time to recurrence for patients whose T_reg_ or T_H_ cells were farther from CE-gland centroids (Fig 8AB; T_reg_ p < 0.01; T_H_ p < 0.05). Fig E in S1 Text further stratifies results by grade: among grade-group 2 cases, distant T_reg_ cells were associated with higher biochemical-recurrence risk (p < 0.01) and T_H_ cells showed a similar, though non-significant, trend (p < 0.1). Grade-group 3 showed comparable patterns but lacked significance (p > 0.2), likely due to smaller sample size. Collectively, these results suggest that immune-cell spatial proximity to CE glands, beyond histologic grade alone, predicts recurrence risk following prostatectomy.

**Fig 8.**
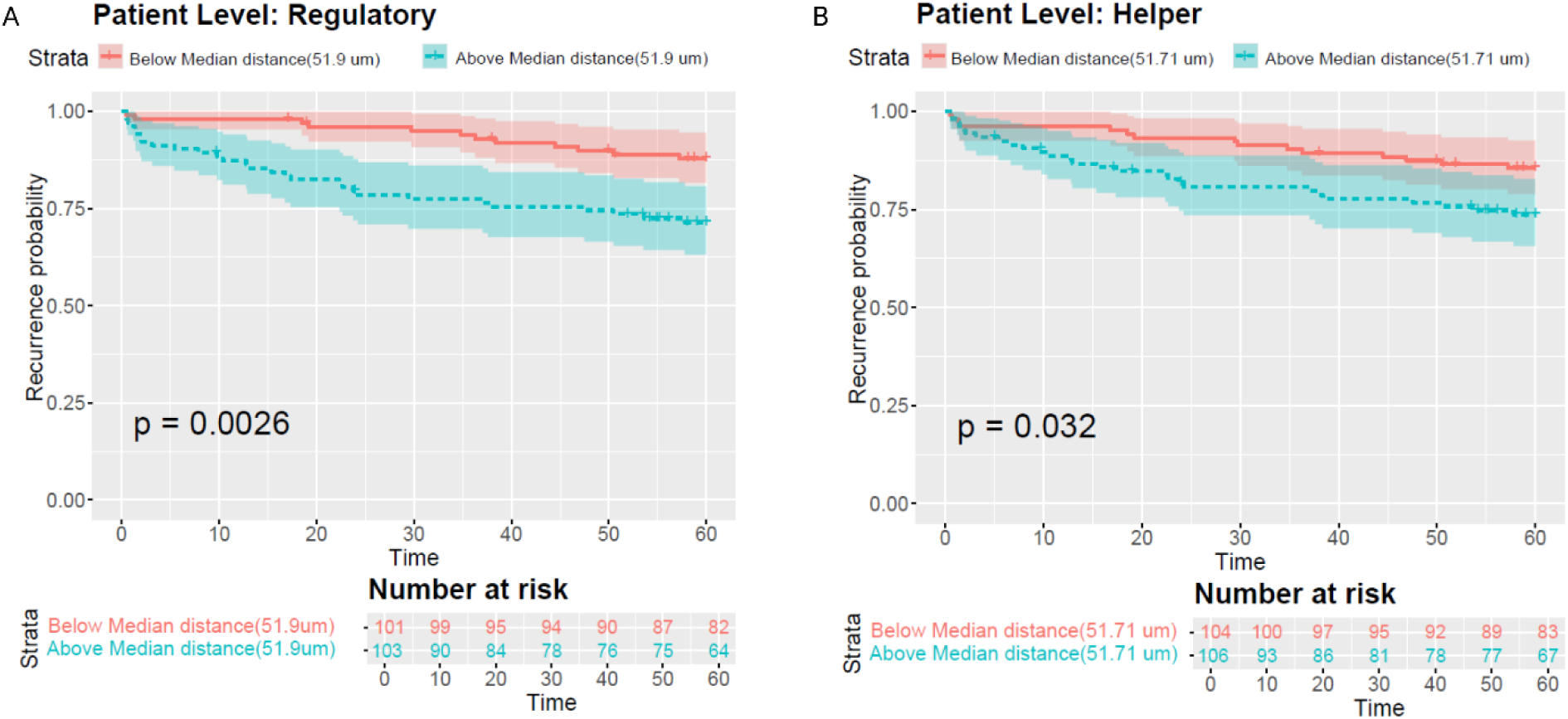
(A) Kaplan–Meier plot of T_reg_-cell distances to nearest CE gland across all grades, dichotomized by median. (B) Kaplan– Meier plot of T_H_-cell distances. Blue = cells farther than median; red = closer than median (T_reg_ p < 0.01; T_H_ p < 0.05).

Using the shape view, we observed grade-related reorganization of MD regions relative to CE glands. Across grades, MD regions enlarged (higher log-mean area, p = 0.07), moved closer to CE glands (smaller boundary-to-boundary distances, p < 0.05), and infiltrated more frequently (distance = 0, p < 0.05; Fig 9). Together, the point- and shape-view analyses highlight macrophage–tumor spatial coupling as a prognostic marker.

**Fig 9.**
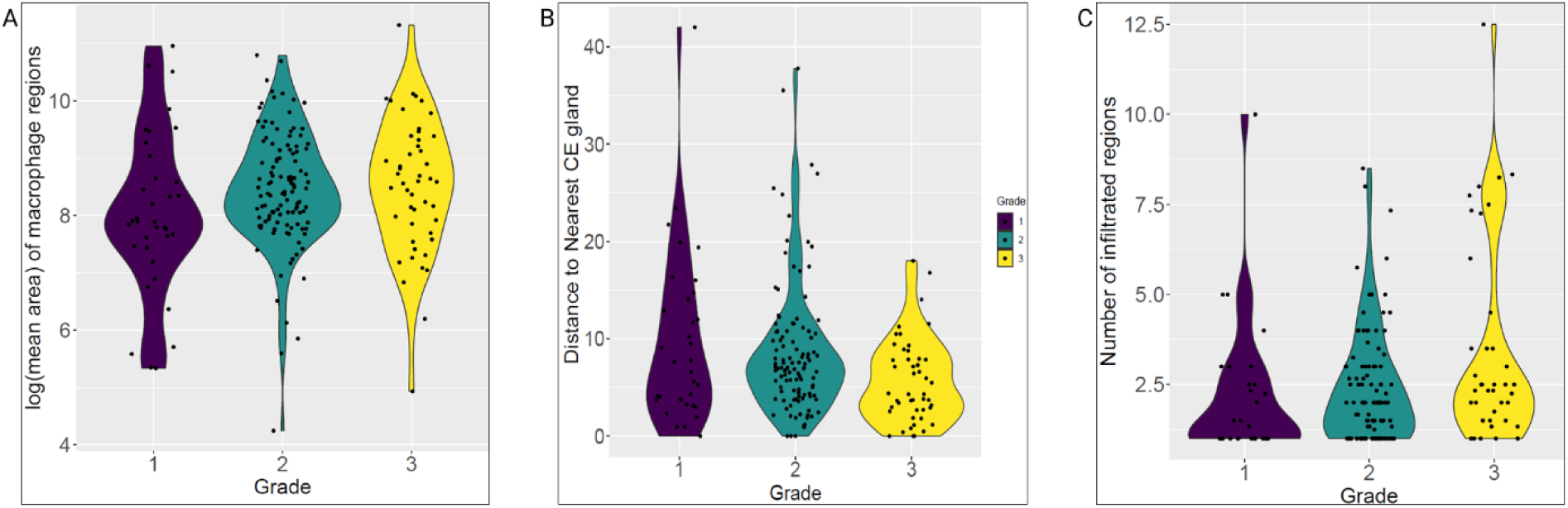
Shape view analysis of macrophage dense (MD) regions. (A) Violin plot of MD-region log-mean area by grade. (B) Violin plot of MD-region distances to nearest CE gland. (C) Number of infiltrating MD regions by grade.

### Software implementation

The cell- and tissue-organization classification model is implemented in TOPAZ (https://chunglab.bmi.osumc.edu/TOPAZ; Fig F in S1 Text), which provides robust visualization and cell- and gland-type classification. Feature extraction and outcome downstream statistical analyses were conducted externally. The TOPAZ workflow consists of five streamlined steps: (1) upload single-cell data (RData or CSV) with corresponding vH&E images and gland masks; (2) select classification biomarkers (p63, CK5) and determine the number of Gaussian components (K = 2 or 3) using histogram visualization; (3) run the GMM to estimate thresholds and apply them to classify cells; (4) overlay results on vH&E images for review; and (5) predict gland shapes using the principal-curve model (Fig 10) and export results.

**Fig 10.**
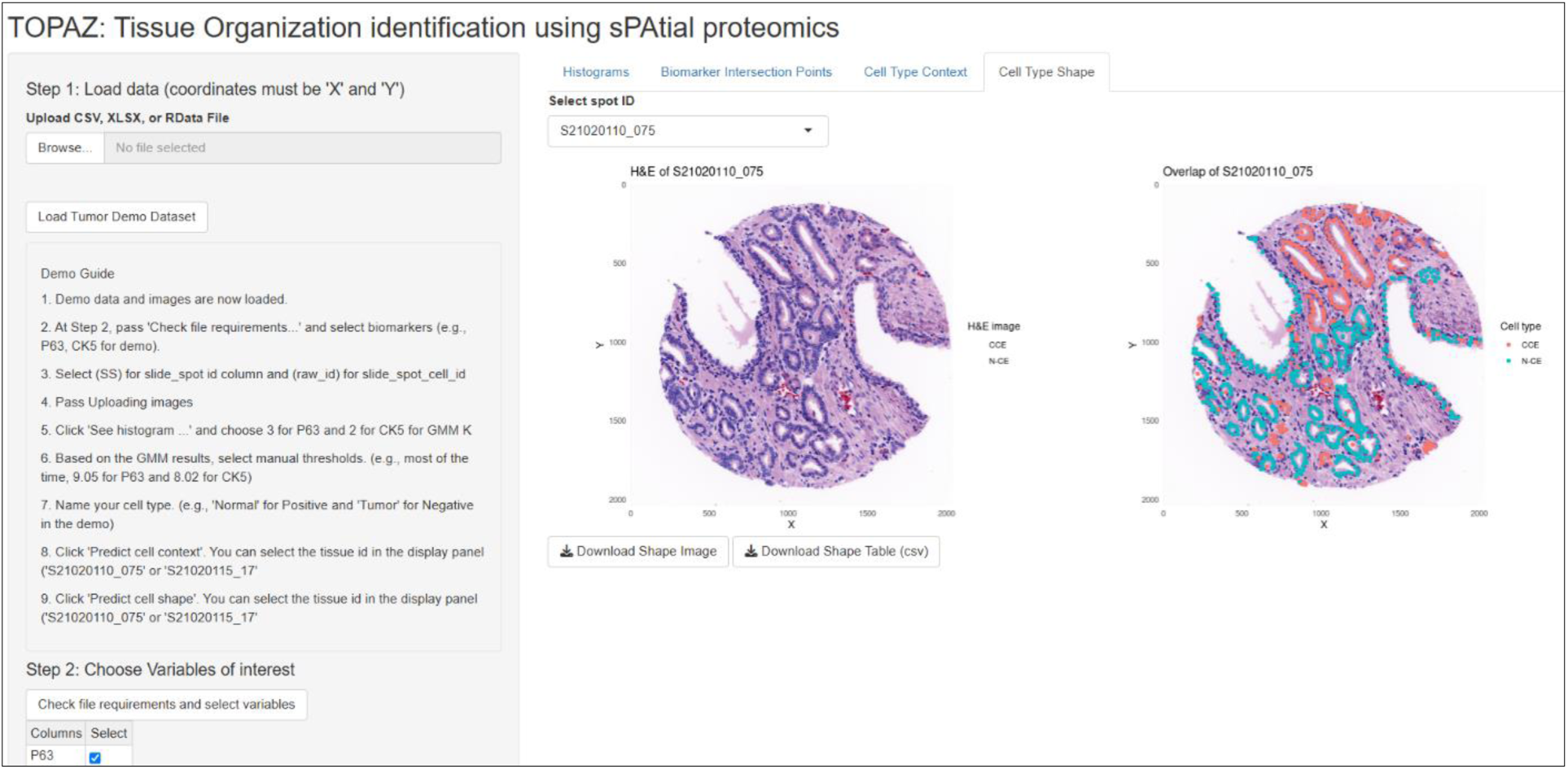
TOPAZ output. Left: vH&E image; Right: principal-curve gland-shape result.

## Conclusion

**TOPAZ** is an annotation-free framework that unifies single-cell identity with gland-level tissue architecture, enabling direct comparison of molecular, geometric, and immunologic features across prostate cancer grades. By integrating multiplexed imaging with probabilistic modeling, the pipeline distinguishes cancerous from non-cancerous epithelial cells (demonstrated by AMACR expression), quantifies macrophage-dense regions and T-cell proximities, and derives spatial features that vary systematically with grade and predict recurrence risk.

Greater T-cell distances and enhanced macrophage–gland interactions—reflected by larger macrophage-dense regions, smaller boundary separations, and more frequent infiltrations—were associated with poorer outcomes. Unlike existing approaches, TOPAZ directly links cellular identity, gland morphology, and immune context to clinical endpoints, providing a comprehensive spatial framework for prostate cancer biology.

The TOPAZ Shiny application delivers these classifications through an interactive interface that visualizes, labels, and exports cell and gland maps, generating datasets suitable for deep-learning model training and validation. Such outputs can accelerate the transition toward fully automated, data-driven digital pathology workflows.

Future improvements of TOPAZ will move beyond automation to enable discovery. Planned developments include adaptive biomarker-threshold learning, seamless integration with clinical platforms such as QuPath, and extension of the principal-curve strategy to a wider range of gland-forming cancers. By merging cell-type classification, gland masks, and spatial statistics within a single, interpretable framework, TOPAZ aspires to serve as a foundation for next-generation spatial pathology—linking multiscale tissue organization to disease trajectory, therapeutic response, and precision oncology.

## Supporting information

**S1 Text. Supplementary information for prostate cancer TOPAZ classification and downstream analyses.**

Additional figures and tables.

## Data Availability

The TOPAZ framework was implemented as a SW and publicly available at https://chunglab.bmi.osumc.edu/TOPAZ. Single cell data and virtual H&E images used for this manuscript are available upon request.

## Funding

Research reported in this publication was supported by the National Cancer Institute of the National Institutes of Health (NIH) under Award Number R01CA249899 (PM). Funder website: https://www.cancer.gov/. The funders had no role in study design, data collection and analysis, decision to publish, or preparation of the manuscript. The content is solely the responsibility of the authors and does not necessarily represent the official views of the National Institutes of Health.

## Competing interests

None declared.

## Author Contributions

KJJ: Conceptualization, Methodology, Software, Formal analysis, Visualization, Writing – Original Draft Preparation.

SG: Methodology, Writing – Review & Editing.

SC: Data curation, Investigation, Writing – Review & Editing.

EM: Investigation, Validation, Writing – Review & Editing.

CC: Investigation, Validation, Writing – Review & Editing.

RW: Writing – Review & Editing.

JDB: Resources, Writing – Review & Editing.

DC: Methodology, Writing – Review & Editing.

FG: Supervision, Writing – Review & Editing.

RM: Supervision, Writing – Review & Editing.

PM: Supervision, Funding acquisition, Writing – Review & Editing.

## Acknowledgements

We thank Yutong Sun for providing valuable feedback on the manuscript during its preparation.

## Notes

### Competing Interest Statement

The authors have declared no competing interest.

